# The separation of Antler Polypeptide and its effects on the proliferation and osteogenetic differentiation of Bone Marrow Mesenchymal Stem Cells

**DOI:** 10.1101/2020.02.17.952499

**Authors:** Wang Ping, Sun Tie-Feng, Li Gang, Zhang Hui-Min, Liu Fan-jie, Gao Zhi-hui, Cao Sheng-nan, Sun Guo-dong, Du Hai-tao, Wang Cong-an, Wang Dan-dan, Shi Bin, Lin Ling

## Abstract

The effects of antler polypeptide on rat bone marrow mesenchymal stem cells (BMSCs) were investigated. Antler polypeptide was separated from Colla Cornus Cervi by ultrafiltration into different samples according to molecular weight: A (molecular weight <800 Da), B (molecular weight 800-1500 Da) and C (molecular weight >1500 Da). The content of antler polypeptide in A, B and C solutions were quantified by high-performance liquid chromatography (HPLC). The effects of antler polypeptide at different concentrations on the proliferation, cell cycle, and osteogenesis of BMSCs were investigated. The highest cell proliferation rate (84.66%) was observed for antler polypeptide B at a concentration of 1.578 × 10^−2^ g/mL. Antler polypeptide B significantly promoted the proliferation of BMSCs with a proliferation index of 38.68%, which was significantly higher than that of the other groups. Antler polypeptide B significantly enhanced the activity of alkaline phosphatase in BMSCs compared to that of blank group (*P* <0.001). Antler polypeptide B increased the BMP7 protein expression in BMSCs. Our data suggested that antler polypeptide may promote the proliferation and osteogenic differentiation of BMSCs.

## Introduction

Antlers have been recorded to have the functions of “invigorating the kidney, nourishing blood, strengthening muscles and bones, promoting blood circulation and detumescence” in *Shen Nong’s Herbal Classic* in traditional Chinese medicine. The annual regenerative cycle of antlers is typical of mammalian-specific cut regeneration and rapid growth, characterized by rapid coordinated regeneration of blood vessels, nerves, cartilage, and bone [1]. Antlers contain a variety of active substances that promote bone cell proliferation and regeneration, suggesting that the active ingredients of antlers could be used to restore and regenerate areas of osteonecrosis [2].

Colla Cornus Cervi, the main extract of antlers, has been used in traditional Chinese medicine for nearly two thousand years. The *Commentaries on the Illustrations* records that “Old deer has good antlers. Boiling antlers to obtain Colla Cornus Cervi, which is better to use as medicine”[3]. Colla Cornus Cervi is rich in protein, accounting for more than 82.49% of all constituents [4]. A previous study revealed that CCC promoted the proliferation and osteogenic differentiation of bone marrow-derived mesenchymal stem cells (BMSCs) [5]. However, the utilization of Colla Cornus Cervi is seriously limited owing to its large molecular weight, which negatively limits its bioavailability [6]. Thus, it is important to improve the bioavailability of Colla Cornus Cervi in order to fully exploit this valuable Chinese medicine resource to better serve human health.

In recent years, increasing researches focused on the physiological functions of peptides. Antler polypeptide, which is isolated from antlers with molecular weight between 0.2 and 10 kDa, have many biological functions and activities [7]. Due to the small molecular weight, simple structures, low immunogenicity, and easily to be absorbed and utilized by cells, antler polypeptide have been widely used clinically to provide nutrition for the body [8-10].

In the present study, we prepared an enzymatic hydrolysate of antler polypeptide by a previously reported method [11], and separated the lysate into three parts according to relative molecular mass by ultrafiltration. Then, high-performance liquid chromatography (HPLC) and the biuret method were used to quantify antler polypeptide in different samples. The effects of different concentrations of the antler polypeptide solutions on the proliferation, cell cycle, and osteogenic differentiation of BMSCs were investigated. Our study lays an experimental foundation for the further development and application of antler polypeptide in medicine.

## Materials and Methods

### Materials

Colla Cornus Cervi is solid glue made from antler tablets by water decoction and concentration. Antler tablets (batch number 150470, Hebei Yabao Pharmaceutical Co. Ltd., China) were identified as originating from red deer by the Chinese Medicine Research Institute of Shandong Academy of Traditional Chinese Medicine. Preparation and identification were carried out according to the 2015 edition of the Chinese Pharmacopoeia section on Colla Cornus Cervi [2].

Pepsin (Meilun Bio, China), Dulbecco’s modified Eagle’s medium-low glucose (DMEM-LG, Hyclone Co. Ltd., USA), 0.25% trypsin-ethylenediamine tetraacetic acid solution (Solarbio Co. Ltd., China), penicillin and streptomycin mixture (Solarbio Co. Ltd., China), phosphate buffered saline (PBS, Solarbio Co. Ltd., China), bovine serum albumin (BSA, purity 98%, Solarbio Co. Ltd., China), fetal bovine serum (FBS, Tianjin Haoyang Biological Products Technology Co. Ltd., China). Cytochrome C (Mr 12,500 Da), aprotinin (Mr 6,500 Da), bacillus enzyme (Mr 1450 Da), glycine-glycine-tyrosine-arginine (Mr 451 Da), glycine-glycine-glycine (Mr 189 Da), and BMSCs were purchased from Saiye Biotechnology (article number RASMX-01001, Guangzhou, China). A rabbit polyclonal antibody against BMP-7 (abcam, Britan), a rabbit polyclonal antibody against GAPDH (SantaCruz, USA), and a horseradish peroxidase (HRP)-conjugated goat anti-rabbit IgG antibody (SantaCruz, USA).

### Instruments

Microfiltration-ultrafiltration-nanofiltration membrane separation tester (BONA-GM-18, Jinan Bona Biotechnology Co. Ltd., China), ZX-002 UV spectrophotometer (Shimadzu, Japan), Waters 2965 HPLC apparatus (Waters Co. Ltd., USA), a KDM-type temperature control electric heating mantle (Hualuyiqi Co. Ltd., China), BP211D Sartorius electronic analytical balance (Germany Sartorius Co. Ltd., Germany), HH-S6 digital constant temperature water bath (Jintan Medical Instrument Factory, China), LGJ-10 freeze dryer (Shengchao Kechuang Biotechnology Co. Ltd., China), −80 °C refrigerator (FOMAS Co. Ltd., USA), plastic culture plates (Corning Co. Ltd.), inverted phase contrast microscope (Olympus Co. Ltd., Japan), CO_2_ incubator (Changsha Huasheng Electronic Technology Co. Ltd., China), microplate reader (Bio-Rad Co. Ltd., USA), ultra-clean workbench (ZHJH-1209, Taiwan Sitea Equipment Co. Ltd., Taiwan).

### Preparation of antler polypeptide

The Colla Cornus Cervi powder and pepsin (4%, m/V) were placed in a 50 mL conical flask and the pH was adjusted to the required level with acetic acid and sodium acetate buffer solutions. The solids were dissolved in water under ultrasonication and enzymolysis in a constant temperature water bath at enzymatic hydrolysis temperature of 40°C. After enzymatic hydrolysis, the enzyme was inactivated by heating to 95 °C for 5 min. The mixture was centrifuged at 7000 g·min^-1^ for 10 min and the supernatant was used to obtain the antler polypeptide hydrolysate.

### Ultrafiltration separation of antler polypeptide

The antler polypeptide hydrolysate was placed in an ultrafiltration system, and ultrafiltration was performed with hollow fiber ultrafiltration membranes with molecular weight cutoffs of 800 and 1500 Da [12, 13]. The ultrafiltration operation temperature was 22-25 °C and the operating pressure was <0.1 MPa. The antler polypeptide was divided as antler polypeptide A (<800 Da), B (800-1500 Da), and C (>1500 Da). The samples were sterilized by cobalt 60 irradiation and stored at −20 °C.

### Determination of antler polypeptide content by biuret method

Standard substance preparation: Standard BSA was prepared and formulated into a standard protein solution of 10.8 mg·mL^-1^ using distilled water. Preparation of biuret reagent: 1.50 g of copper sulfate (CuSO_4_·5H_2_O) and 6.0 g of sodium potassium tartrate (KNaC_4_H_4_O_6_·4H_2_O) were dissolved in 500 mL distilled water, added 300 mL of 10% NaOH solution, and diluted to 1 000 mL with distilled water. Standard curve: took 12 tubes into two groups, added 0, 0.2, 0.4, 0.6, 0.8, 1.0 mL standard BSA solution, complemented to 1 mL with distilled water, then added 4 mL of biuret reagent and placed at room temperature (20∼25 °C) for 30 min. Absorbance value at 540 nm was determination with a microplate reader. Taking the concentration as the abscissa and the absorbance as the ordinate, the standard curve was obtained: y=0.0464x-0.0022, *R*^2^=0.9998. The absorbance of antler polypeptide samples was determined using the same method, and each sample was performed in triplicate. According to the standard curve, the antler polypeptide content was calculated as follows: 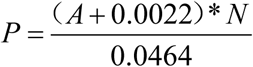, *A*: absorbance value, *N*: the dilution rate, *P*: the content of antler polypeptide.

### Determination of antler polypeptide content by HPLC

The reference substances (cytochrome C, aprotinin, bacillus enzyme, glycine-glycine-tyrosine-arginine, and glycine-glycine-glycine) were accurately weighed. The mobile phase was configured using 0.1% (mass fraction) peptide standard solutions of different relative molecular weights. Sample preparation was performed using 200 μL of antler polypeptide A, B, or C, which was diluted with pure water 10-times, filtered with a microporous membrane (0.2-0.5 μm polytetrafluoroethylene or nylon filter membrane). The injection volume was 10 μL. A TSKgel G2000 SWXL (300 mm × 7.8 mm) column was used for separation with a mobile phase of acetonitrile/water/trifluoroacetic acid (45:55:0.1, *v/v/v*). The UV detection wavelength was 220 nm, the flow rate was 0.5 mL·min^-1^, and the column temperature was 30 °C.

To validate the method, its accuracy (six injections), repeatability (six mixed reference analyses), and stability (0, 2, 4, 6, 12, 24 h) were evaluated. The relative standard deviations (RSDs) of the determined peak areas were less than 3%, confirming the viability of the method. The chromatograms of the samples were analyzed by reference to the peak times of the five standard substances. Then, using gel-permeation chromatography (GPC) data processing software, the chromatographic data of the sample was substituted into the calibration curve equation to calculate the relative molecular weight of the peptide in the sample and its distribution range. The sum of the relative peak area percentages of the antler polypeptides was calculated by peak area normalization.

### Culture of BMSCs

BMSCs were placed in a 25-cm^2^ flask and cultured in complex medium at 37 °C in a 5% CO_2_ incubator. The complex medium was DMEM-LG containing 20% FBS, 100 U/mL penicillin and 100 µg·mL^-1^ streptomycin. The complex medium was changed every 3 days. The fourth generation of BMSCs were collected and washed three times with PBS. After digestion with 0.25% trypsin, single cell suspension was harvested for further experiment. BMSCs were identified by flow cytometry (CD29, CD44 and CD90 were positive with the expression rates >70%, whereas CD34, CD45 and CD11b were negative with the expression rates <5%).

### Determination of proliferation of BMSCs

After digestion, the BMSCs cell concentration was adjusted to 2 × 10^4^ mL^-1^. Cells were seeded in a 96-well plate at a density of 2×10^3^ cells per well and cultured at 37 °C in a 5% CO_2_ incubator for 12 h. The supernatant was discarded and 50 μL of antler polypeptide A, B, or C and 50 μL medium were added, while 50 μL PBS and 50 μL medium were used as a blank group. After culturing for 24 h, 48 h, or 72 h, the supernatant was discarded and 100 μL PBS and 10 μL of 5 mg·mL^-1^ 3-(4,5-dimethylthiazol-2-yl)-2,5-diphenyltetrazolium bromide (MTT) was added to each well. The plate was then incubated at 37 °C in a 5 % CO_2_ incubator for 4 h. Then, the supernatant was removed, 150 μL DMSO was added to each well. The plate was oscillated for 10 min and absorbance value at a wavelength of 490 nm was detected using a microplate reader. The effects of antler polypeptide A, B, and C on the proliferation rate of BMSCs were calculated as follows: proliferation rate = average OD value of experimental group / average OD value of control group × 100%.

### Cell cycle assay

Flow cytometry (propidium iodide staining) was used to investigate the BMSCs cell cycle. BMSCs were seeded into six-well plates at a density of 2×10^6^ per well, which were randomly divided into blank group, Colla Cornus Cervi group, and antler polypeptide A group, antler polypeptide B group, and antler polypeptide C group. After culturing for 24 h, BMSCs were digested with trypsin and resuspended in ice-cold PBS at a density of 3×10^6^ mL^-1^ and incubated with RNase in a 37°C water bath for 30 min. The cells were subsequently incubated with propidium iodide at 4°C in the dark for 30-60 min and analyzed using a flow cytometer. The cell cycle of BMSCs was analyzed using software of ModFit Lt for mac V1.01.

### Detection the activity of alkaline phosphatase (ALP)

The BMSCs cells were seeded in a six-well plate at a density of 5×10^5^ per well, and divided into blank group, Colla Cornus Cervi group, antler polypeptide A group, antler polypeptide B group, and antler polypeptide C group. After culturing for 48 h, the supernatant was discarded. Then, 250 μL of lysate was added into the six-well plate and the cells were lysed for 30 min. The absorbance at 520 nm was detected using the ALP detection kit according to the manufacturer’s instruction. Three independent experiments were performed.

### Western blot analysis

The cells of each group were lysed in RIPA lysis buffer at 4°C for 15 min. The lysates were cleared by centrifugation (12,000 rpm) at 4°C for 30 min to collect total protein. About 20 μg protein samples were then separated by electrophoresis in a 10% SDS (sodium dodecyl sulfate) polyacrylamide gel and the transferred onto a polyvinylidene fluoride membrane. After blocking the non-specific binding sites for 60 min with 5% non-fat milk, the membranes were incubated with a rabbit polyclonal antibody against BMP-7 (at 1:1000 dilution) at 4°C overnight. The membranes were then washed with TBST (tris-buffered saline with tween-20) three times at room temperature for 15 min. After washing, the target protein was probed with the horseradish peroxidase (HRP)-conjugated goat anti-rabbit IgG antibody (at 1: 2000 dilution) at 37°C for 1 h. After three washes, the membranes were developed by an enhanced chemiluminescence system. The protein levels were normalized with respect to GAPDH protein level which was detected using a rabbit polyclonal antibody against GAPDH (at 1: 5000 dilution).

### Statistical analysis

Statistical processing was performed using SPSS 19.0. The experimental data are expressed as 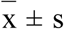. The least significant difference method was used for comparison between groups. *P*<0.05 was considered statistically significant.

## Results

### Ultrafiltration separation of antler polypeptide

After ultrafiltration, three different colored antler polypeptide liquids were obtained as shown in Figure1. The solution of the antler polypeptide A (molecular weight <800 Da) was clear and transparent, antler polypeptide B (molecular weight 800-1500 Da) was light yellow, sticky, no obvious odor, and antler polypeptide C (molecular weight >1500 Da) was black brown.

**Figure 1.**
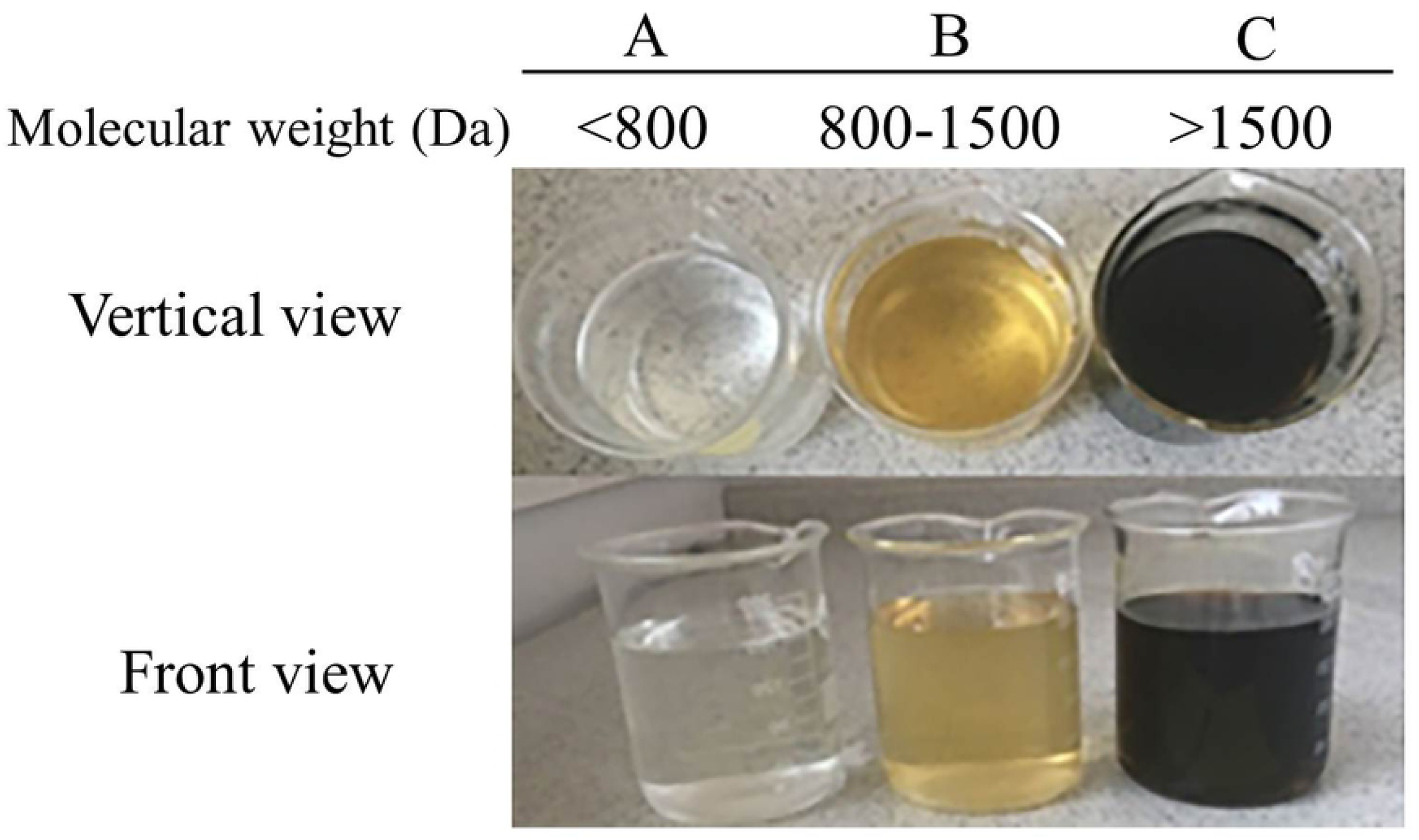
Three different colored antler polypeptide liquids with different molecular weight separated through ultrafiltration. (A) Antler polypeptide A, molecular weight <800 Da. (B) Antler polypeptide B, molecular weight 800-1500 Da. (C) Antler polypeptide C, molecular weight >1500 Da.

### Determination of antler polypeptides by biuret method

The concentration of A, B and C samples was 6.319 mg/mL, 7.181 mg/mL, and 7.776 mg/mL, respectively. The peptide content of A, B and C samples determined by biuret method was 0.602 mg/mL, 8.976 mg/mL, and 38.88 mg/mL, respectively. The results were shown in table1.

**Table 1.** The peptide content of A, B and C samples determined by biuret method

| samples | volume | average<br>absorbance | concentration<br>( mg/mL ) | antler polypeptide content<br>( mg/mL ) |
| --- | --- | --- | --- | --- |
| A | 0.7mL | 0.291 | 6.319 | 0.602 |
| B | 0.8mL | 0.331 | 7.181 | 8.976 |
| C | 0.2mL | 0.358 | 7.776 | 38.88 |

Note: Antler polypeptide A (molecular weight <800 Da) sample was concentrated as 15 times.

### HPLC analysis of antler polypeptide

The results were shown in Figure 2. The antler polypeptides with molecular weights <800 Da, 800-1500 Da, and >1500 Da were successfully separated. Comparing the retention times of the five standard materials (cytochrome C, aprotinin, bacillus enzyme, glycine-glycine-tyrosine-arginine, and glycine-glycine-glycine), the peptide peak (peak area 933.80927) with the molecular weight 800-1500 Da eluted at 14.279∼15.351 min showed that the content of antler polypeptide was significantly higher than that of the other samples (Table 2).

**Table 2.** The peak area results of different samples detected by HPLC (n=3)

| Group | Eluted time/min | Peak area |
| --- | --- | --- |
| A | -- | -- |
| B | 14.460 | 933.80927 |
| C | 14.907 | 20.30075 |

Note: The peptide peak of different samples with the molecular weight 800-1500 Da eluted at 14.279~15.351 min.

**Figure 2.**
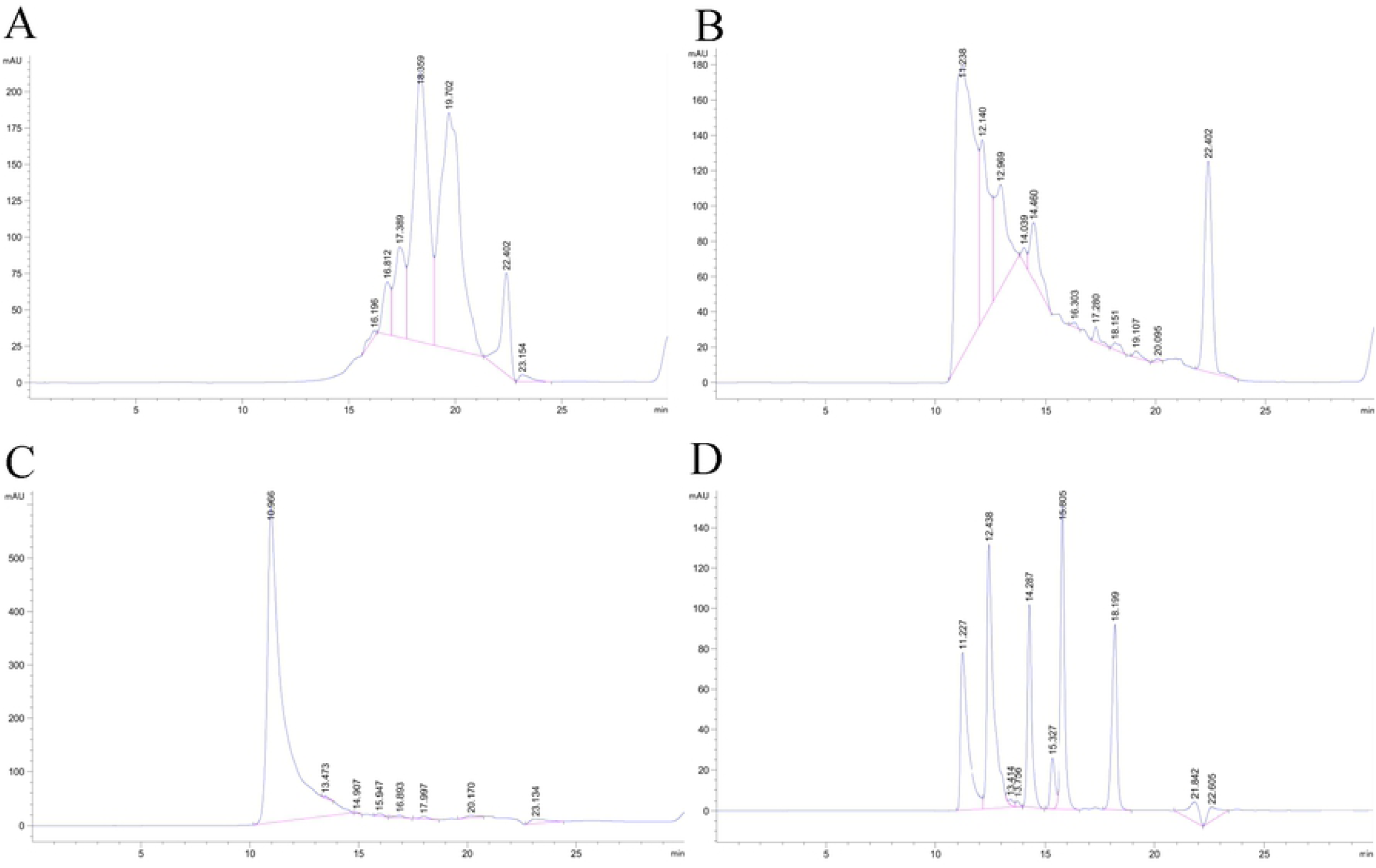
HPLC chromatograms of antler polypeptides solutions. Antler polypeptide A, molecular weight <800 Da; Antler polypeptide B, molecular weight 800-1500 Da; Antler polypeptide C, molecular weight >1500 Da; Peptide standard samples, 1, cytochrome C, 2, aprotinin, 3, bacillus enzyme, 4, glycine-glycine-tyrosine-arginine, 5, glycine-glycine-glycine. Elution time, 14.461-15.454 min.

### Morphological observation of BMSCs

The morphology of the BMSCs was observed under an inverted phase contrast microscope (Figure 3). After two days of cell growth, the BMSCs showed single or multiple cell clones. On the day 5, the BMSCs covered the bottom with a density of 90% and presented long wedge shapes.

**Figure 3.**
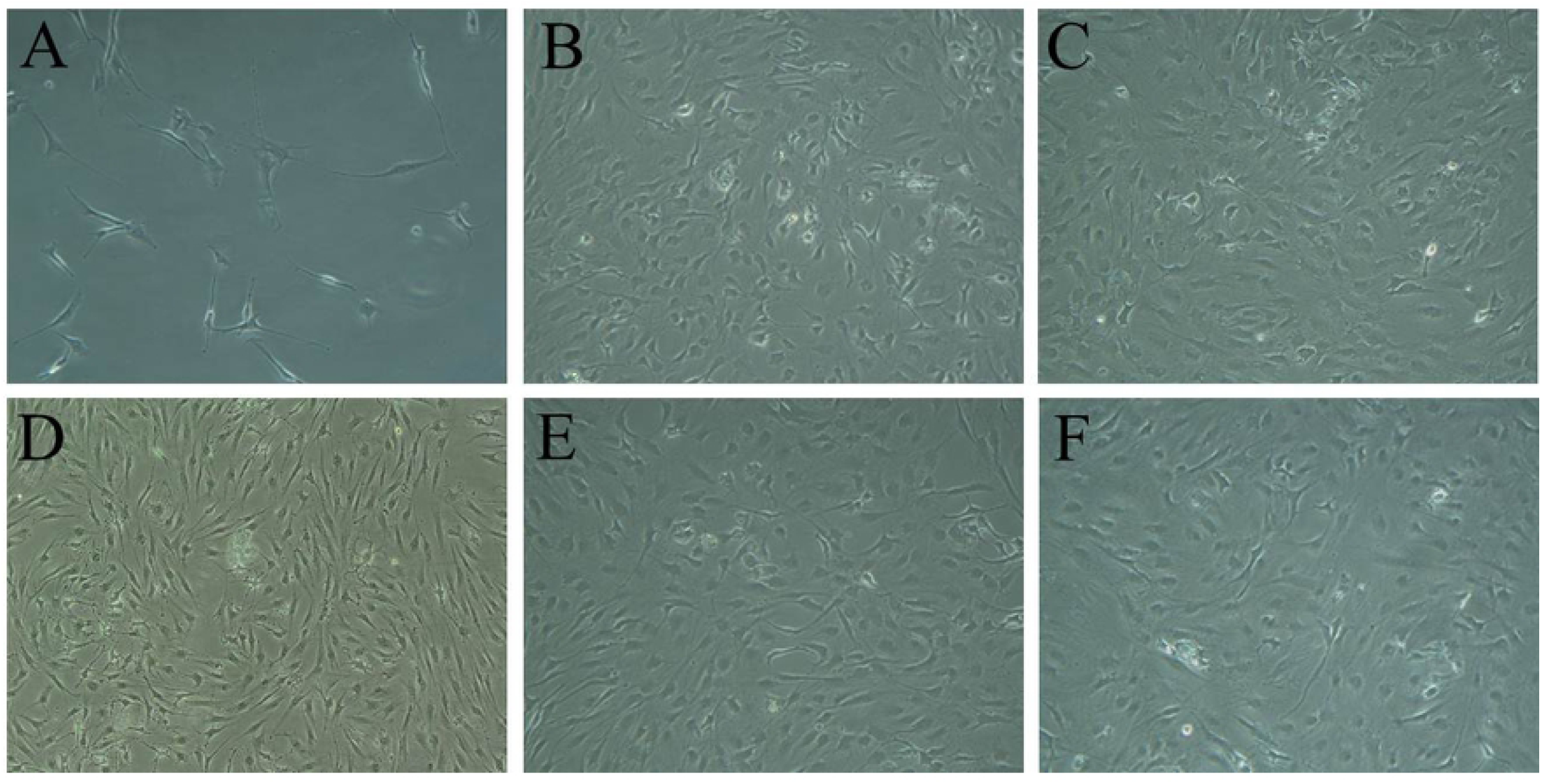
BMSCs observed under an inverted phase contrast microscope. (A) BMSCs cultured with complex medium for 2 d. (B) BMSCs cultured with complex medium for 5 d. (C) BMSCs cultured with complex medium containing antler polypeptide A for 5 d. (D) BMSCs cultured with complex medium containing antler polypeptide B for 5 d. (E) BMSCs cultured with complex medium containing antler polypeptide C for 5 d. (F) BMSCs cultured with complex medium containing Colla Cornus Cervi for 5 d. Magnification, ×200.

### Effects of antler polypeptide on the proliferation of BMSCs

BMSCs were cultured with complex medium containing different concentrations of antler polypeptides A, B, C, or Colla Cornus Cervi respectively to evaluate the proliferation of BMSCs. The results were shown in table 3. The antler polypeptides A, C and Colla Cornus Cervi group have no obvious effect on the proliferation of BMSCs. Antler polypeptide B significantly promoted BMSCs proliferation, and the effect was most positive when concentration of antler polypeptide B was 1.578 × 10^−2^ g/mL. As the concentration decreased, the proliferation rate of BMSCs decreased.

**Table 3.**
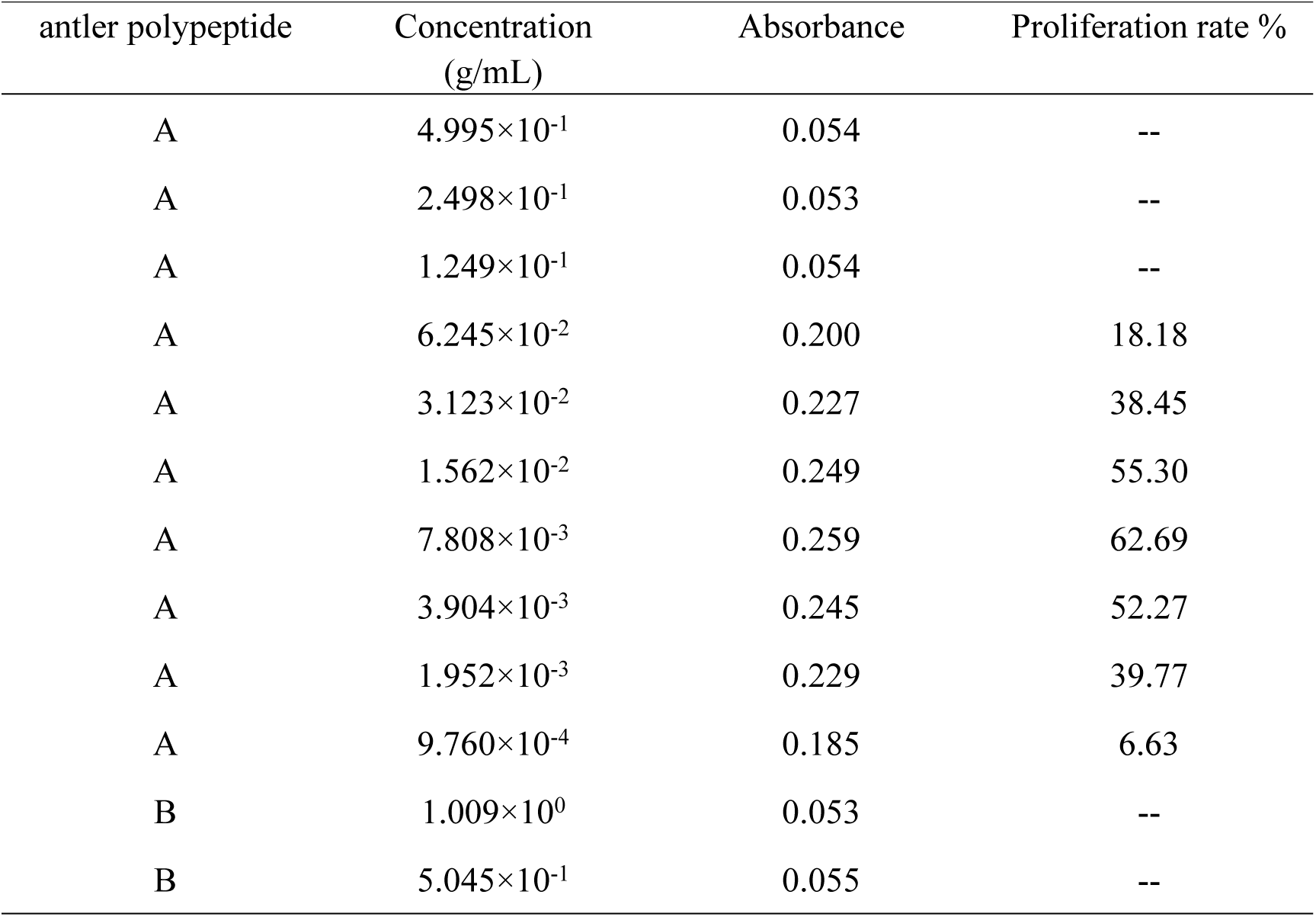
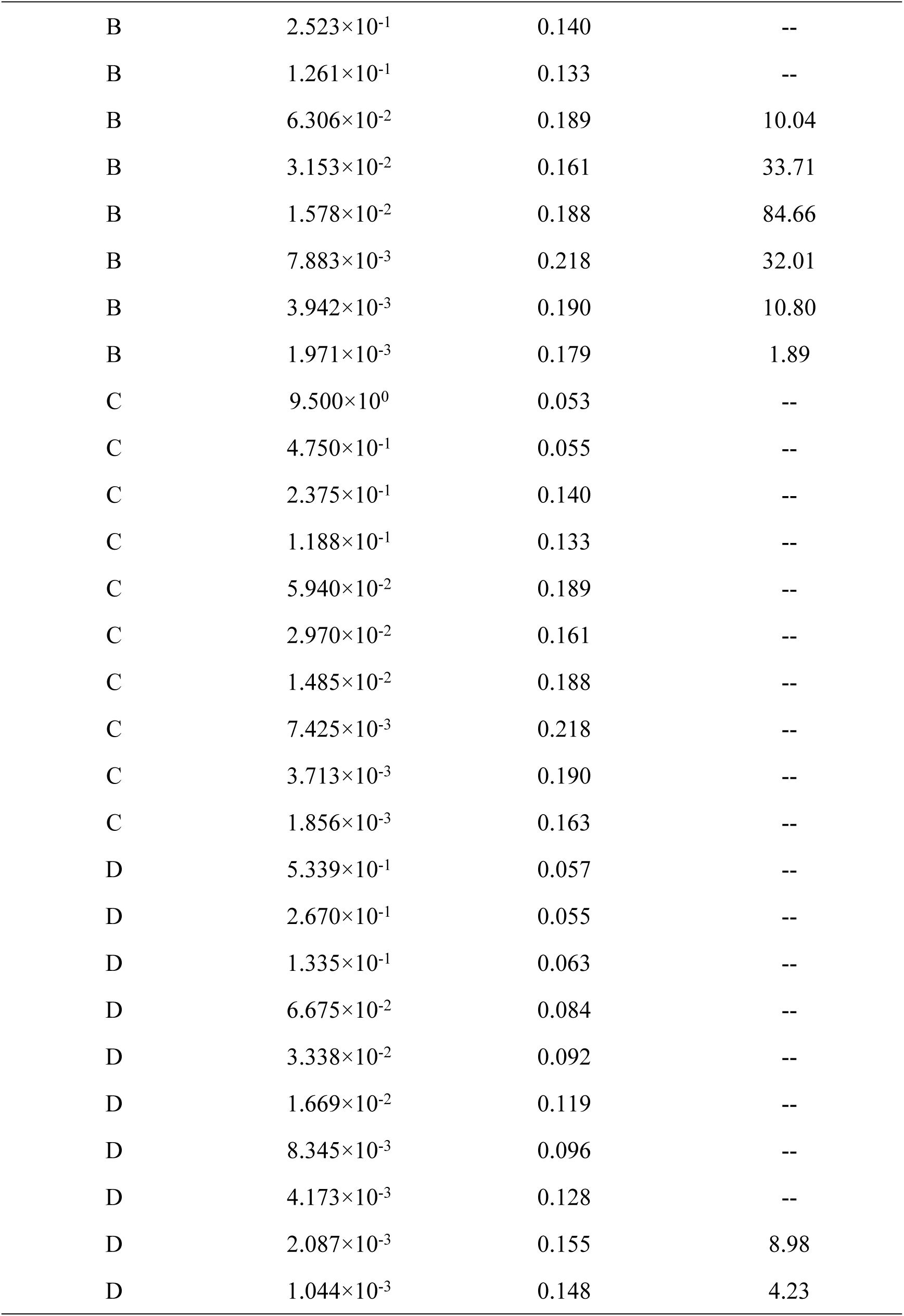
Effects of different concentrations of antler polypeptide on the proliferation of BMSCs (n=4)

| antler polypeptide | Concentration<br>(g/mL) | Absorbance | Proliferation rate % |
| --- | --- | --- | --- |
| A | $4.995 \times 10^{-1}$ | 0.054 | -- |
| A | $2.498 \times 10^{-1}$ | 0.053 | -- |
| A | $1.249 \times 10^{-1}$ | 0.054 | -- |
| A | $6.245 \times 10^{-2}$ | 0.200 | 18.18 |
| A | $3.123 \times 10^{-2}$ | 0.227 | 38.45 |
| A | $1.562 \times 10^{-2}$ | 0.249 | 55.30 |
| A | $7.808 \times 10^{-3}$ | 0.259 | 62.69 |
| A | $3.904 \times 10^{-3}$ | 0.245 | 52.27 |
| A | $1.952 \times 10^{-3}$ | 0.229 | 39.77 |
| A | $9.760 \times 10^{-4}$ | 0.185 | 6.63 |
| B | $1.009 \times 10^0$ | 0.053 | -- |
| B | $5.045 \times 10^{-1}$ | 0.055 | -- |
| B | $2.523 \times 10^{-1}$ | 0.140 | -- |
| B | $1.261 \times 10^{-1}$ | 0.133 | -- |
| B | $6.306 \times 10^{-2}$ | 0.189 | 10.04 |
| B | $3.153 \times 10^{-2}$ | 0.161 | 33.71 |
| B | $1.578 \times 10^{-2}$ | 0.188 | 84.66 |
| B | $7.883 \times 10^{-3}$ | 0.218 | 32.01 |
| B | $3.942 \times 10^{-3}$ | 0.190 | 10.80 |
| B | $1.971 \times 10^{-3}$ | 0.179 | 1.89 |
| C | $9.500 \times 10^0$ | 0.053 | -- |
| C | $4.750 \times 10^{-1}$ | 0.055 | -- |
| C | $2.375 \times 10^{-1}$ | 0.140 | -- |
| C | $1.188 \times 10^{-1}$ | 0.133 | -- |
| C | $5.940 \times 10^{-2}$ | 0.189 | -- |
| C | $2.970 \times 10^{-2}$ | 0.161 | -- |
| C | $1.485 \times 10^{-2}$ | 0.188 | -- |
| C | $7.425 \times 10^{-3}$ | 0.218 | -- |
| C | $3.713 \times 10^{-3}$ | 0.190 | -- |
| C | $1.856 \times 10^{-3}$ | 0.163 | -- |
| D | $5.339 \times 10^{-1}$ | 0.057 | -- |
| D | $2.670 \times 10^{-1}$ | 0.055 | -- |
| D | $1.335 \times 10^{-1}$ | 0.063 | -- |
| D | $6.675 \times 10^{-2}$ | 0.084 | -- |
| D | $3.338 \times 10^{-2}$ | 0.092 | -- |
| D | $1.669 \times 10^{-2}$ | 0.119 | -- |
| D | $8.345 \times 10^{-3}$ | 0.096 | -- |
| D | $4.173 \times 10^{-3}$ | 0.128 | -- |
| D | $2.087 \times 10^{-3}$ | 0.155 | 8.98 |
| D | $1.044 \times 10^{-3}$ | 0.148 | 4.23 |

### Effects of antler polypeptide B on the growth of BMSCs

When BMSCs were cultured with complex medium containing antler polypeptide B, the proliferation rate of BMSCs was highest at 48 h and slower after 72 h (Figure 4A). The proliferative effect for both these time-points was significantly better than that at 24 h. Compared with the blank group, antler polypeptide B significantly enhanced BMSCs proliferation at 24 h, 48 h, and 72 h (*P* <0.01). Thus, antler polypeptide B promoted the growth of BMSCs.

**Figure 4.**
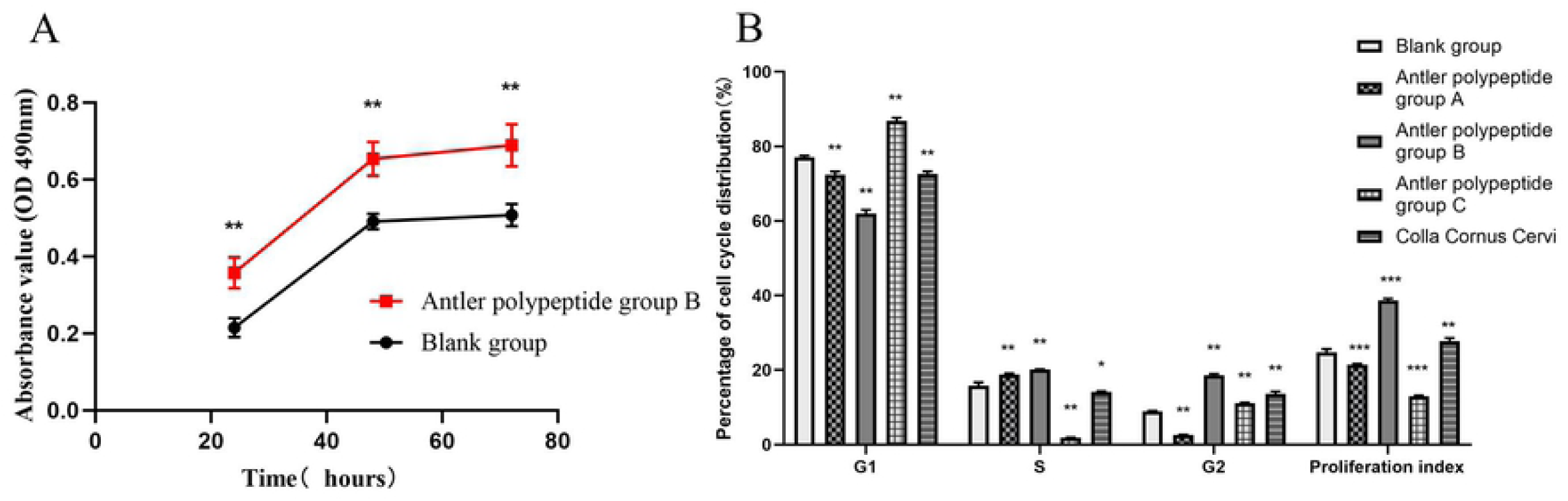
Antler polypeptide B promoted the growth of BMSCs. (A) Cell proliferation assay showing that antler polypeptide B promoted the proliferation of BMSCs. (B) Effects of different antler polypeptide groups on the cell cycle of BMSCs. The proliferation index reflected cell cycle process and proliferative ability of BMSCs. Antler polypeptide B group showed a significantly higher proliferation index. * P<0.05, ** P<0.01, *** P<0.001, compared with blank control group.

### Effects of different groups of antler polypeptides on the cell cycle of BMSCs

The proliferation index reflected cell cycle process and proliferative ability of BMSCs. The proliferation index of each group was as follows (Figure 4B): antler polypeptide B group (38.68%), Colla Cornus Cervi group (27.74%), blank group (24.70%), antler polypeptide A group (21.39%) and antler polypeptide C group (12.99%). Compared with the blank group, antler polypeptide B group showed a significantly higher proliferation index, indicating that antler polypeptide B with a molecular weight of 800-1500 had the strongest proliferative effect on BMSCs.

### Effect of antler polypeptide on Alkaline phosphatase activity of BMSCs

The activity of ALP was significantly increased in antler polypeptide B group compared with blank control group (*P* < 0.001, Figure 5A). The Colla Cornus Cervi group, antler polypeptide groups A and C had significant effects on the ALP activity compared with blank control group (*P* < 0.05). The results indicated that antler polypeptide can enhance the activity of ALP in BMSCs.

**Figure 5.**
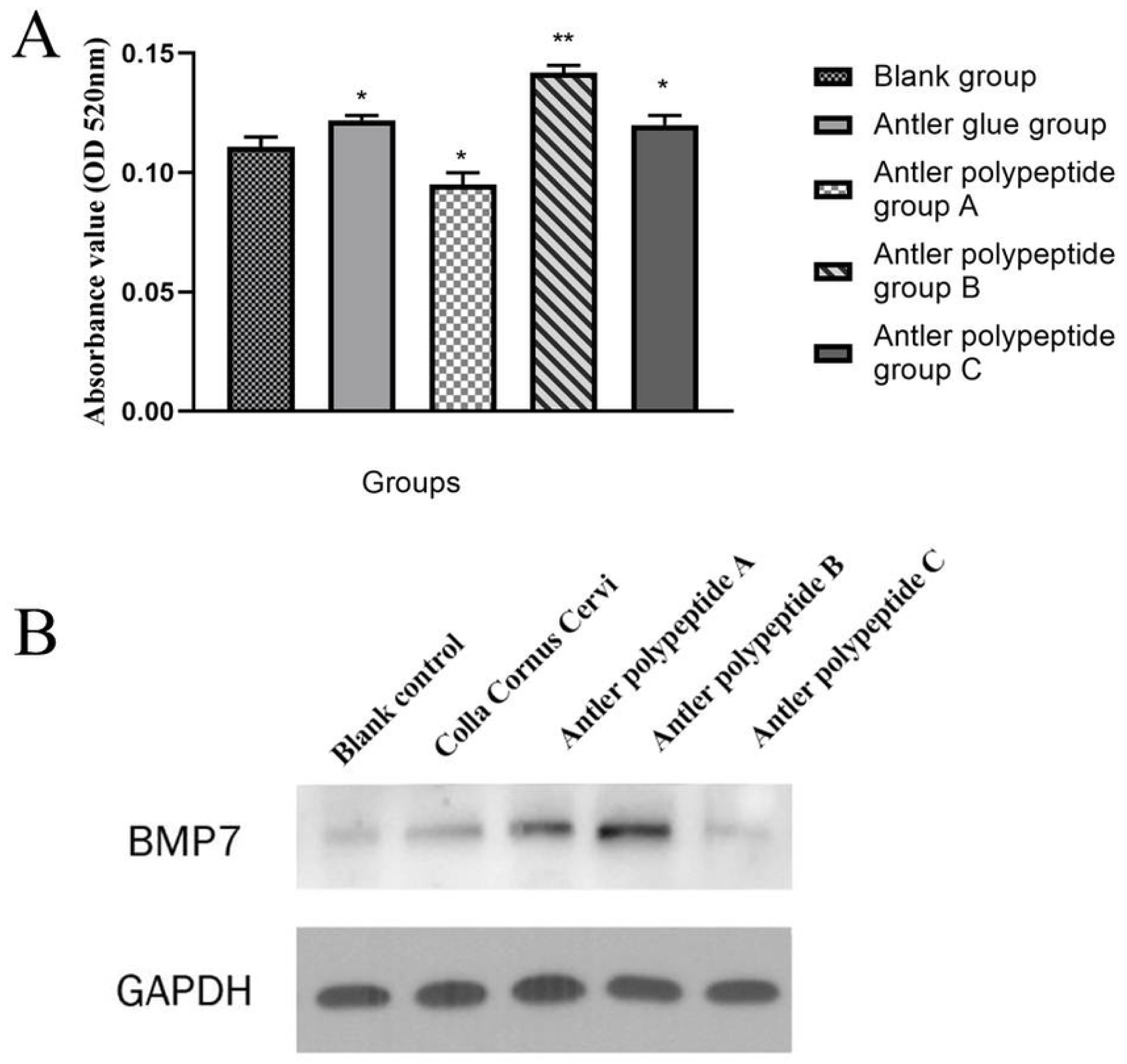
Effect of antler polypeptide on osteogenic differentiation of BMSCs. (A) Antler polypeptide enhanced the ALP activity of BMSCs. (B) Antler polypeptide promoted BMP7 protein expression in BMSCs. * P<0.05, ** P<0.01, compared with blank control group.

### Effect of antler polypeptide on BMP7 expression

Western blotting results showed a BMP7 band at the expected size. BMP7 protein expression was markedly increased in the antler polypeptide A, B and Colla Cornus Cervi group compared with blank control group (Figure 5B). The antler polypeptide B group showed maximum BMP7 expression. The results indicated that antler polypeptide enhanced the BMP7 expression in BMSCs.

## Discussion

Antler polypeptide which are separated from antlers have been reported to have biological activity in different diseases [7]. Antler polypeptide play important roles in antioxidant and anti-fatigue effects. Antler polypeptide improved the proliferation ability of mouse spleen lymphocytes and enhanced the immune function [14]. Antler polypeptide inhibited bone loss and prevented osteoporosis [9]. Xie et al [15] reported that velvet antler polypeptide partially rescue facet joint osteoarthritis.

In the present study, antler polypeptide were extracted from Colla Cornus Cervi powder and separated as antler polypeptide A (<800 Da), B (800-1500 Da) and C (>1500 Da) group according to the molecular weights. Antler polypeptide were identified through HPLC, and the content of antler polypeptide in group B was significantly higher than that of the other samples. The effects of antler polypeptide on biological functions of BMSCs were further investigated. Results confirmed that antler polypeptide B with a molecular weight of 800-1500 Da significantly promoted growth and proliferation of BMSCs, and the optimum concentration was 1.578×10^−2^ g/mL. Cell cycle analysis revealed that antler polypeptide B significantly increased the proliferation index of BMSCs. These results suggested that antler polypeptide promoted BMSCs growth and proliferation through promoting the cell cycle process of BMSCs.

The effect of antler polypeptide on osteogenic differentiation of BMSC was further investigated. Results showed that antler polypeptide B markedly enhanced ALP activity, which is an important indicator of osteogenic differentiation. Furthermore, antler polypeptide B significantly enhanced BMP7 protein expression in BMSCs. BMP7, also known as bone morphogenetic protein 7, is a member of the transforming growth factor β family. BMP7 has many biological functions, including regulating cell growth, proliferation, differentiation, apoptosis, and inducing bone formation. Previous study revealed that overexpression of BMP7 significantly promoted the osteogenic differentiation of BMSCs [5]. Kang et al [16] found that BMP7 enhanced osteogenic differentiation of BMSCs transfected with a BMP7-recombinant-vector. BMP7 enhanced the osteogenic differentiation of human dermal-derived CD105^+^ fibroblast cells through the Smad and MAPK pathways [17]. These researches combined with our results indicated that, BMP7 may associate with osteogenic differentiation of BMSCs promoted by antler polypeptide.

For the mechanism study, it was reported that antler polypeptide may regulate OPG/RANKL/RANK signaling pathway to prevent osteoporosis [18]. Antler polypeptide enhanced the differentiation of BMSCs and inhibited the growth of osteoclasts by regulating NF-κB signaling pathway [19]. Antler polypeptide inhibited the activity of IL-1 and IL-6, promoted the proliferation of BMSCs and inhibited bone loss [15]. Ren et al [6] reported that pilose antler aqueous extract promoted the proliferation and osteogenic differentiation of bone marrow mesenchymal stem cells by stimulating the BMP-2/Smad1 and 5/Runx2 signaling pathway. Our present study suggested that antler polypeptide promoted proliferation and osteogenic differentiation of BMSCs. In further studies, the molecular mechanisms of antler polypeptide on proliferation and osteogenic differentiation of BMSCs will be investigated.

In conclusion, antler polypeptide was extracted from Colla Cornus Cervi powder and confirmed through HPLC analysis. Antler polypeptide (molecular weight 800-1500 Da) significantly promoted the proliferation of BMSCs with a proliferation index of 38.68%. Antler polypeptide increased the activity of alkaline phosphatase and enhanced BMP7 protein expression in BMSCs. Our study suggested that antler polypeptide promoted the proliferation and osteogenic differentiation of BMSCs. The present research lays an experimental foundation for the further development and application of antler polypeptide in medicine.

## Acknowledgements

This work was supported by grants from the Major Science and Technology Innovation Projects of Shandong Province (2018CXGC1310), the Shandong Key Research and Development Plan (2017CXGC1306; 2016ZDJS07A23; 2016GSF2020420), the China Postdoctoral Science Foundation (2019M662420), the Natural Foundation of Shandong Province (ZR2015HM061; ZR2019MH134), the Distinguished Experts of Taishan Scholar Project (ts201511074), National training Program for innovative backbone talents of traditional Chinese Medicine, TCM Technology Development Project of Shandong Province (2019-0543), Academic promotion plan of Shandong First Medical University (2019QL003) and Science and Technology Plan of Shandong Academy of Medical Sciences (2018-23). The funders had no role in study design, data collection and analysis, decision to publish, or preparation of the manuscript.

